# A computational model for learning from repeated trauma

**DOI:** 10.1101/659425

**Authors:** Alfred P. Kaye, Alex C. Kwan, Kerry J. Ressler, John H. Krystal

**Affiliations:** Yale University Department of Psychiatry, New Haven, CT; VA National Center for PTSD Clinical Neuroscience Division, West Haven, CT; Yale University Department of Neuroscience, New Haven, CT; McLean Hospital, Division of Depression and Anxiety Disorder, Belmont, MA; Harvard Medical School, Department of Psychiatry, Boston, MA

## Abstract

Traumatic events can lead to lifelong inflexible adaptations in threat perception and behavior which characterize posttraumatic stress disorder (PTSD). This process involves associations between sensory cues and internal states of threat and then generalization of the threat responses to previously neutral cues. However, most formulations neglect adaptations to threat that are not specific to those associations. In order to incorporate non-associative responses to threat, we propose a computational theory of PTSD based on adaptation to the frequency of traumatic events using a reinforcement learning momentum model. Recent threat prediction errors generate momentum that influences subsequent threat perception in novel contexts. This model fits data acquired from a mouse model of PTSD, in which unpredictable footshocks in one context accelerate threat learning in a novel context. The theory is also consistent with epidemiological data showing that PTSD incidence increases with the number of traumatic events, as well as the disproportionate impact of early life trauma. Since the theory proposes that PTSD relates to the average of recent threat prediction errors rather than the strength of a specific association, it makes novel predictions for the treatment of PTSD.

## Introduction

Computational psychiatry seeks to define psychiatric disorders in terms of fundamental algorithms for survival rather than only as pathological states (1-3). Quantitative models may allow personalization of mental health care, insight into the nature of the disorder, inform neurobiological investigations into psychiatric disorders, or predict the trajectory of symptoms (4-6). For example, depression has been conceived as an adaptation to periods of low reward availability (7). Similarly, hallucinations have been conceptualized as resulting from excessive weighting of prior expectations for auditory stimuli in a Bayesian model (8-9). One approach to describing a computational function of a neural system is using David Marr’s three levels of analysis (10) (Figure 1A), which seeks to map connections between computational goals, algorithmic procedures to achieve them, and the neurobiological substrate underlying these processes.

**Figure 1.**
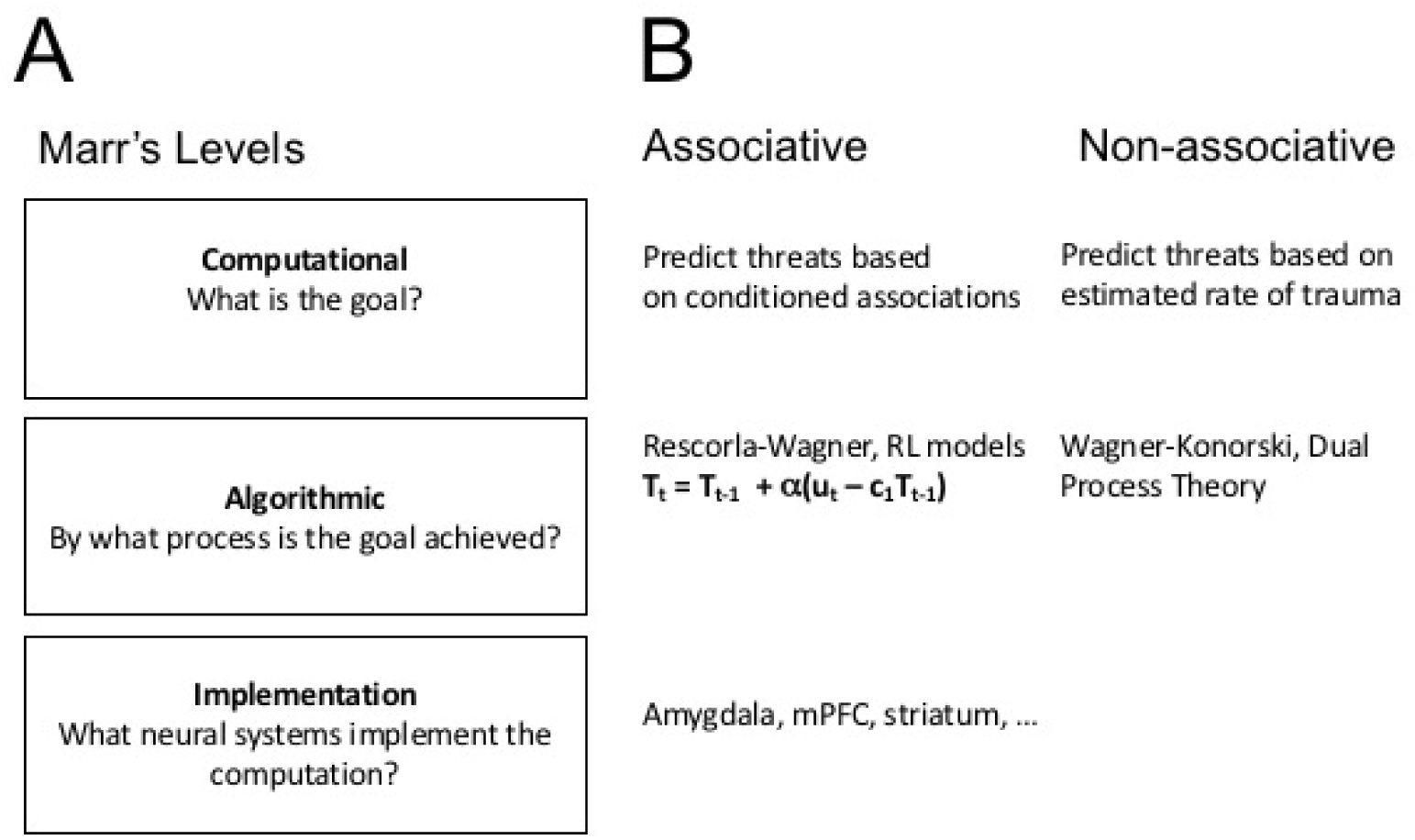
David Marr’s Levels of Analysis for computational neuroscience as applied to PTSD. (A) Definition of the three levels of analysis from ref. 7. (B) Application of those levels to associative learning (left) and non-associative learning (right) in PTSD. (left) Associative learning is a well-characterized system with a clear computational goal of ethological relevance (Computational), a mathematically defined formal model (Algorithm), and neural circuit mechanisms (Implementation). (right) Non-associative learning is less well-understood. The goal posited here is that it’s purpose is to predict threats based on repetition of traumatic events (Computational). Schematized models exist (Algorithmic) but lack a formal mathematical model, and the neurobiological correlates of this are not fully understood (Implementation).

Posttraumatic stress disorder has a computational description that organizes theory and neurobiological data across Marr’s three levels - associative fear learning (Figure 1B, refs. 11-15). Learning models have been successfully applied to PTSD and underlie current conceptualizations of the disorder and treatment options (16-17). PTSD is seen as an extreme outcome of associative fear learning, which in turn is a fundamental mechanism for predicting threats based on previous experience (18). In this model, PTSD occurs when life-threatening situations create potent associations between sensory reminders of the traumatic event and the emotional experience of fear (17). The intensity of this association then motivates a person to avoid (19) future trauma cues, limits extinction of the fear memory (20), and supports the subsequent formation of new fear memories via generalization and second-order conditioning (21). This process can be described mathematically, enabling learning parameters to be precisely measured during new associative learning in a laboratory setting (18). The precision with which associative learning can be controlled has enabled neurobiological studies into circuit mechanisms in both humans and animals (22).

In contrast, non-associative learning – increases (sensitization) or decreases (habituation) in response to a repeated stimulus (23) – is a prominent component of PTSD that lacks a formal algorithmic description (Figure 1B). In humans, repeated traumatic events increase the probability of developing PTSD and may change the nature of the disorder (24-25). Core PTSD symptoms, such as hyperarousal, inherently involve an exaggerated response to sensory cues – importantly, these cues need not be associated with the traumatic event to trigger the response (26) but may instead result from sensitization of neuromodulatory systems (27-28). Neurobiological studies in animals have shown that stress enhances both innate defensive behaviors (29) and learning about unrelated fear cues (30). There are conceptual models of how habituation and sensitization occur (Dual Process Theory, ref. 31; Wagner-Koniorsky Theory, ref. 32), which center the role of arousal in changing the response to a stimulus with repetition. However, these models lack the algorithmic detail and clear relation to survival value of Rescorla-Wagner and related reinforcement learning (RL) models (33). This has limited the ability to parametrically manipulate and therefore understand non-associative learning in PTSD patients and animal models.

Here, we posit an ecological role for non-associative learning in estimating the frequency of predator attacks (or other violence). We then apply a Bayesian approach to understand how well an ideal agent could estimate predation risk from its own life history. We show that a natural consequence of this approach is that early life trauma has disproportionate impact on estimated risk even when controlling for the number of traumatic events. After describing the behavior of such an ideal Bayesian agent, we turn to a recently developed RL model (7, 34-36) in order to integrate associative and non-associative learning. Non-associative learning becomes more important as traumas occur in more different contexts and less distant times. The RL model points towards novel interventions and future neurobiological approaches to improve PTSD symptoms.

## Methods

### Models of threat estimation

Two models of threat estimation are identified and compared: (1) a Bayesian model, in which an agent experiences events (attacks) and attempts to estimate the frequency of those attacks and (2) a reinforcement learning agent, which experiences events (attacks) in contexts (all attacks occur in different contexts) over time and must estimate the threat in each environement. The reinforcement learning model is then compared with behavioral data for a mouse undergoing a stress procedure.

### Bayesian attack model

At each time step, events (attacks) are binomially distributed with probability of attack *p*_*a*_ for 700 time steps (Figure 2a). Deaths occur with probability *p*_*d*_ contingent on an attack occurring. The agent’s estimate of *p*_*a*,_ and *p*_*d*_ is derived from the sequence of attack observations (***x***_**1**_ = 0,0,1 … 0) according to Bayes’ rule

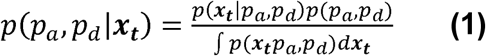

using a Markov Chain Monte Carlo sampler with a flat prior at time t=0. Specifically, an affine invariant ensemble MCMC sampler (MCMC Hammer, ref. 37) toolbox for Matlab with 31 walkers was used to estimate the posterior. For subsequent timepoints, Bayesian estimation is performed with the prior distribution as the posterior of the previous time step.

**Figure 2.**
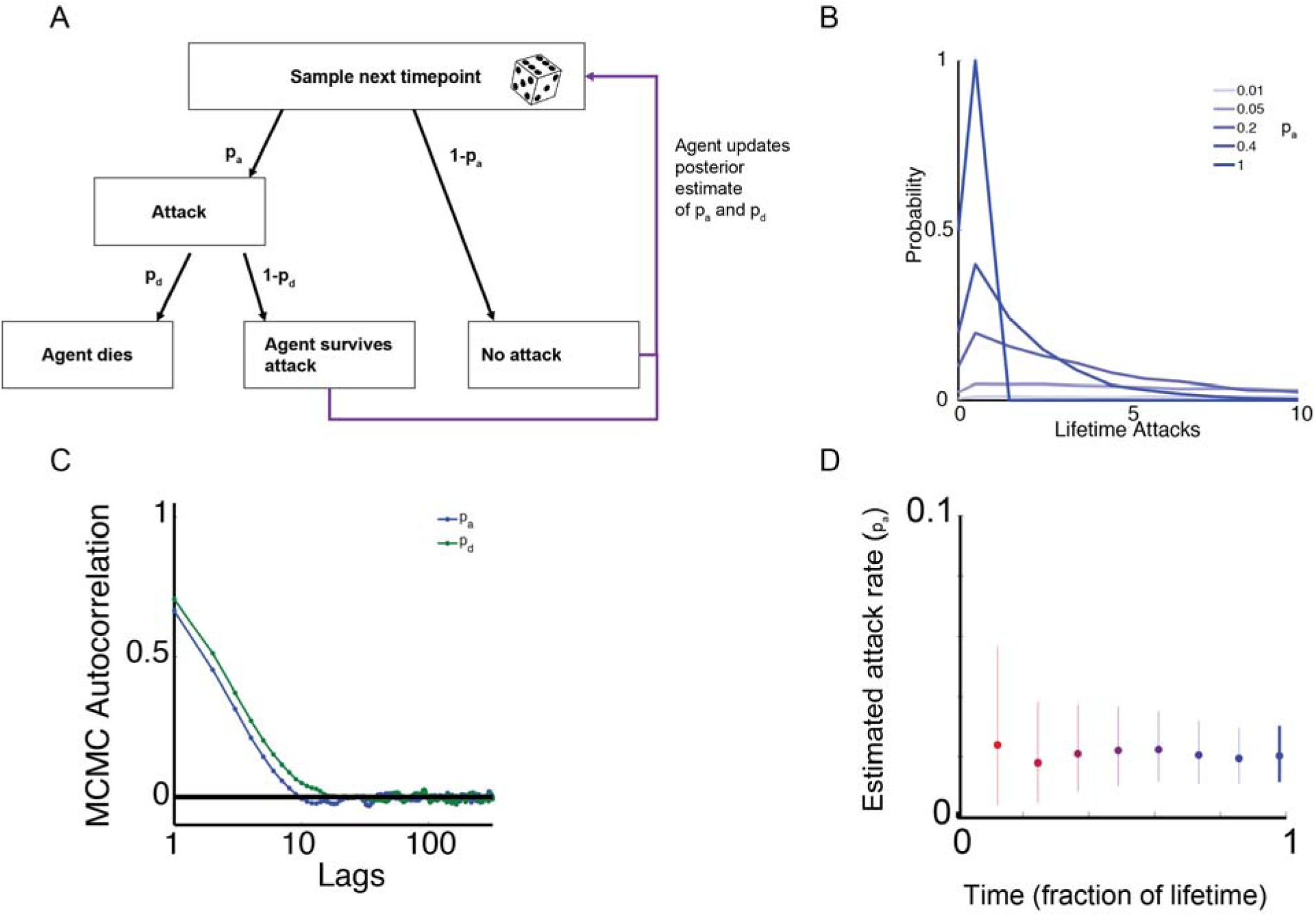
A Bayesian observer can measure the rate of traumatic attacks. (A) Schematic of a simplified doubly stochastic model of attacks (i.e., traumatic events). Attacks occur raxsndomly at each timepoint with a fixed probability *p*_*a*_. Conditional on attacks occurring, agents die with probability *p*_*d*_. If agents survive, they estimate the ongoing probability of attacks according to Bayes’ rule. (B) Agents must estimate *p*_*a*_ and *p*_*d*_ in an information-poor environment. The number of attacks experienced by the typical agent is low, usually 3-5 over the course of a lifetime for *p*_*a*_=0.2, a typical value for the lethality of predator attacks (30). (C) The Bayesian estimator of the posterior estimates of *p*_*a*_ and *p*_*d*_ for a typical example sequence of attacks (*p*_*a*_ = 0.01, *p*_*d*_ = 0.2) shows convergence for *p*_*a*_ (blue)and for *p*_*d*_ (green). The autocorrelation of the MCMC sampler goes to zero rapidly at long timelags for both parameters, demonstrating convergence in the MCMC sampler. (D) As the agent continues over its lifetime (red to blue map), the estimate of *p*_*a*_ slowly narrows (vertical lines, 95% intervals). Greater time allows the agent to accumulate greater evidence about the true value of *p*_*a*_.

Autocorrelated attack rate time series were generated for an AR(1) autoregressive process

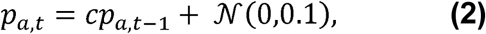

where *p*_*a,t*_ is the attack rate at time t, *c* is a constant equal to the correlation of successive time steps, and *𝒩*(*μ,σ*) is normally distributed noise with *μ* mean (and standard deviation *σ*. Simulations used the arima function in Matlab. N=10,000 simulated lifetime attack rate time series were generated, then for each an agent’s experienced attack time series was generated and the MCMC Hammer estimator was then used to progressively estimate attack rates as above.

### Reinforcement learning models

In temporal difference learning, threat at time t in context c (*T*_*c,t*_) is learned from a sequence of unconditioned stimuli (*u*_*t*_) which produce prediction errors according to

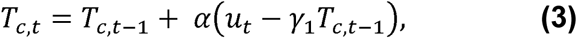

where *α* is a learning rate and *γ*_l_ is a decay rate constant. Equation 3 is referred to as RL model in the Results section, and describes the formation of associative threat learning. The addition of a momentum term (7) allows prediction errors from different states to influence one another according to an RL momentum model

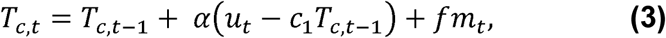

where f is a scaling constant and *m*_*t*_ is the momentum at time t. This momentum term is defined by

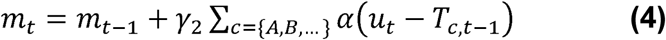

in which the sum of decayed prediction errors across all contexts *c* = {*A, B*,….} with momentum decay constant *γ*_2_. This can lead to either oscillatory behavior or slow summation of prediction errors across states depending on *γ*_2_. Reinforcement learning models (RL – equation 3, RL with momentum – equation 4) were fit to smoothed freezing (sliding window, 15s) on days 1, 6, and 7. Inputs to the model were shock times and threat was fit for both Context A and Context B. Parameters for each model were fit using maximum likelihood estimation in Matlab. Maximum likelihood fit was compared by calculating the Bayes Information Criterion (BIC) for RL and RL with momentum models at the single animal level for both stressed and unstressed mice.

### Stress enhanced fear learning

All procedures were carried out in accordance with the ethical guidelines of the National Institutes of Health and were approved by the Institutional Animal Care & Use Committee of Yale University. 8-12 week old C57Bl/6 male mice were stressed using using the Stress-Enhanced Fear Learning model (30), which has been shown to lead to long-lasting enhancement of fear and anxiety behaviors in both mice (30) and rats (38). This model consists of 15 unpredictable footshocks (1mA, 1s) with random intershock intervals between 4 and 8 minutes. For contextual fear experiments, a second context (Context B) was used on day 6, in a separate room with different ambient auditory, visual, tactile, and olfactory characteristics. On Day 6, a single 1mA 1s shock was administered after 5 minutes, and then freezing was assessed for 5 more minutes. On day 7, mice were returned to Context B for 10 minutes. MedAssociates boxes were used for all footshock experiments, and freezing was assessed as complete cessation of movement other than breathing (motion <18 a.u.) with automated VideoFreeze software.

## Results

Previous approaches to computational modeling of PTSD have focused on defining changes in associative learning after traumatic experience (11-15). PTSD is thus framed as a consequence of underlying mechanisms for predicting threat based on previous *associations*. In contrast, we were interested in whether PTSD might arise from an agent estimating the *frequency* of threat exposure. In order to determine how an ideal observer would estimate the frequency of threat exposure, we first posit a simplified model of exposure to repeated traumatic events. By constructing an ideal Bayesian observer of these traumatic events, we establish a baseline for what can be inferred from repeated events without association. We then turn to a recently developed reinforcement learning model (7, 34-36) to integrate non-associative learning (about the frequency of threat) with associative learning (about the associations of threat). Finally, we fit the reinforcement learning model to data derived from mice undergoing Stress-enhanced Fear Learning (SEFL), a rodent model of PTSD (30). We then consider the implications of our findings for treatment and future research into the neurobiology of PTSD.

## Model 1 – PTSD as trauma rate estimation

An organism must estimate the threat of violence to adapt to it. This process of estimation must necessarily involve information gathered across timescales, since threat may increase suddenly or may increase over long periods (39). Longer timescale estimation of threat involves integrating experience in disparate environments.

To consider a concrete example: predator attacks are events which carry a significant probability of death (20% for mice exposed to an owl, ref. 40). If the probability of death is high, then the animal will experience few attacks before dying (Figure 2B). In this information-poor environment, the animal must maximize the available information in estimating the rate of such attacks. In order to determine how well an ideal observer could do under such conditions, we constructed a simple probabilistic model with a fixed probability of attacks *p*_*a*,_ and probability of dying per attack *p*_*d*_ at each time point (Figure 2A). Using a Markov Chain Monte Carlo sampler, we were able to estimate the posterior distribution of *p*_*a*_ (Figure 2C), which makes it possible to identify the estimate available to a Bayesian observer. As expected, variance in *p*_*a*_ decreases progressively over the lifetime of the agent as more samples become available (Figure 2D).

The disproportionate impact of early life stress (ELS) on adult behavior (39) is explained by the Bayesian trauma rate model. Childhood traumatic experiences have a strong impact on adult brain structure and function (41). Life History Theory explains this by positing that stressful experiences in childhood provide information about organismal strategies that will be adaptive in the adult environment (42). We evaluated the Bayesian trauma rate estimator in two scenarios with the same total number of traumatic events, one in which traumas occur early in life (ELS) and one in which they are spread across the lifespan (Figure 3A). Variance in 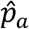 decreases with time in both models, as traumatic events reduce uncertainty in the true rate of violence (Figure 3B). However, over the course of the lifespan the ELS model shows a higher estimated rate of violence 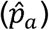. Thus, the increased response to ELS does not require specialized critical period mechanisms, but instead arises naturally in a normative estimator of violence rate.

**Figure 3.**
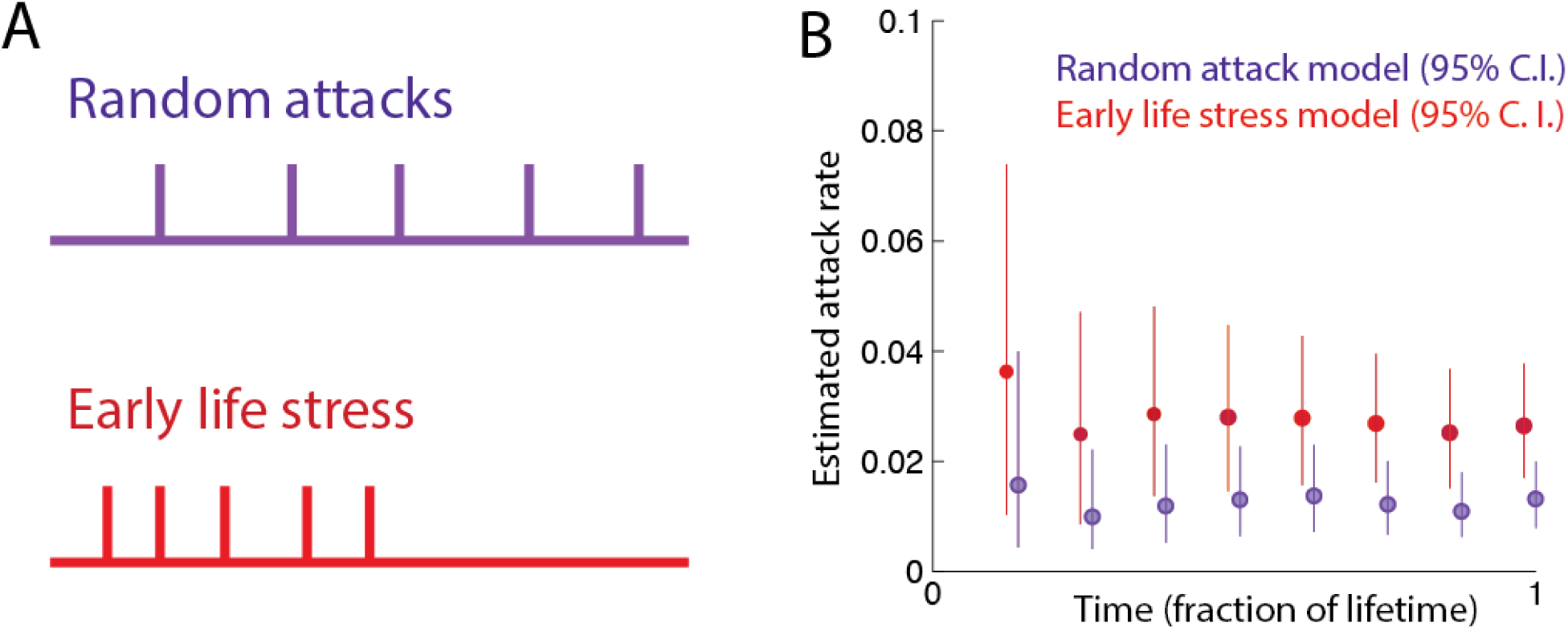
Early life traumas have a disproportionate effect on the estimated attack rate. (A) Characteristic examples of two distributions of attack frequencies. In the random attack model, attacks are uniformly distributed across the lifespan. In the early life stress (ELS) model, an identical number of attacks are uniformly distributed across the first half of the lifespan. (B) Bayesian agents’ posterior distributions for attack rate sequentially measured across the lifespan, for the random and early attack models (*p*_*a*_= 0.01 and *p*_*d*_ = 0.2). The discrepancy between estimated and true attack rate is greatest at the start of life due to a higher density of attacks in the early life stress model. Over the course of the lifespan, these two models arrive at similar estimates.

## Model 2 – PTSD as threat momentum

Normative Bayesian models can explain the performance of an ideal behavior, but are difficult to implement in biological systems due to the computational difficulty in integrating probability distributions to find the posterior (43). It can therefore be useful to define more biologically plausible models which can then be compared to the performance of the ideal Bayesian observer (37). Reinforcement learning (RL) is a flexible class of models that can be used to learn in real time from experience. Unlike Bayesian models, RL involves updating stored values of stimuli or actions based on set learning rules. Parameters of RL models can then be fit to empirical behavioral data of animals or human subjects, to derive differences in parameters between groups. RL models can also be used to explain learning processes, or to identify neural processes that map onto learning processes.

In this section, we propose that a recently proposed RL momentum model (7, 34-36) can explain features of PTSD not explained by classical associative learning models. Traumatic events may come in clusters, so learning from trauma involves combining information from distinct experiences that occur close in time. The momentum model as applied to neuropsychiatric disorders suggests that a common tendency, or mood, may underlie motivated behaviors over a period of time. For intuition into the reason why traumatic events occur together, consider an agent subject to predation risk. Empirical measurements of predator-prey interactions confirm the existence of large fluctuations in predator number (39), which are also predicted by mathematical models of predator-prey interactions such as the Lotka-Volterra equations. In order to adapt to time-varying predator rates, an organism must be capable of tracking the rate of attacks it experiences.

Classical RL models, such as temporal difference learning (Figure 4A), enable an organism to associate threatening experiences with the context in which they are experienced. However, threats in one context do not influence threats in another (Figure 4A). In contrast, in the RL-momentum model, traumatic events occurring close in time but in unrelated environments contribute to a slowly varying momentum term (Figure 4B), which can be thought of as a pervasive mood biasing subsequent experience. Momentum carries information about recent threats, allowing the agent to correctly assess risk in a changing environment. The ideal length of time for momentum to persist depends on how long threats persist (Figure 4C). When attacks are uncorrelated in time, there is no advantage to momentum and the optimal momentum learning rate (highest correlation to the underlying threat rate) is zero, reducing the RL momentum model to a classical RL model. When attacks are correlated (Figure 4C, light blue), a substantial improvement in threat estimation can be obtained by including the momentum parameter. The long-time scale of optimal threat adaptation offers a potential explanation for the persistence of PTSD symptoms. If threat momentum, rather than the specific association with the initial traumatic event, were the source of PTSD symptoms, then this would have substantial implications for the understanding of PTSD.

**Figure 4.**
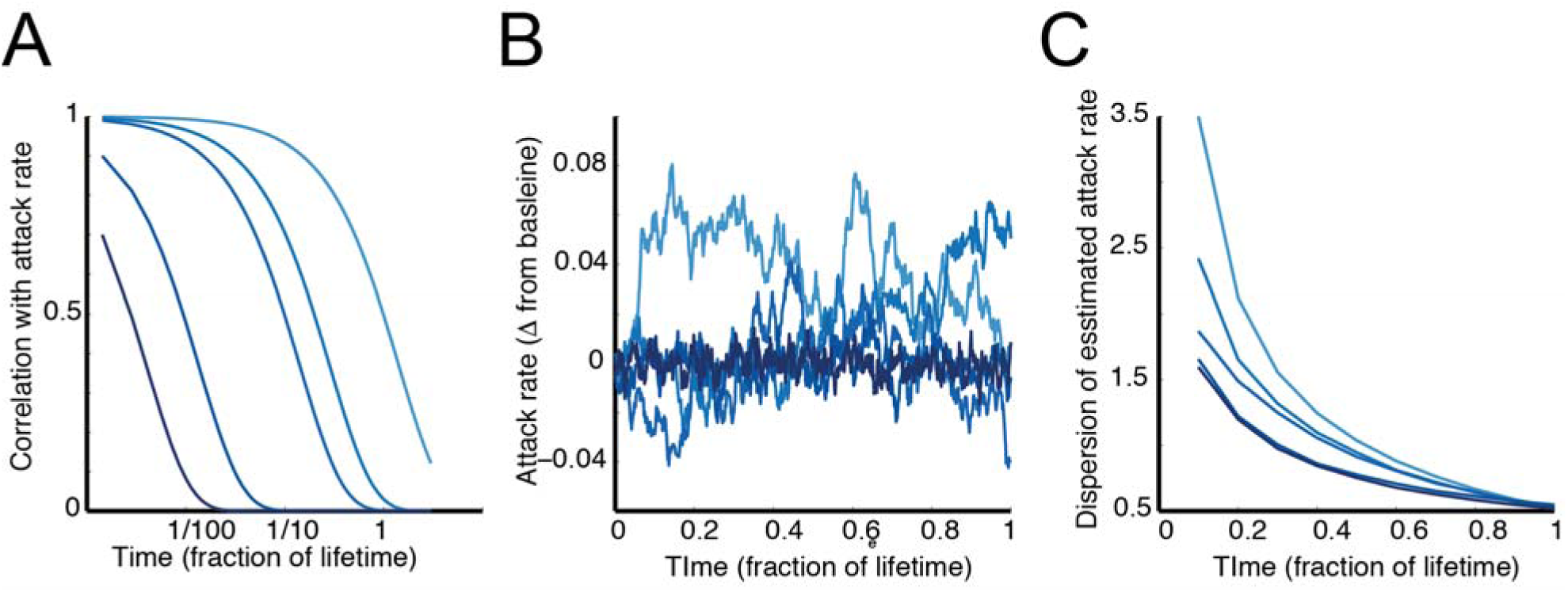
Varying attack rates lead to misestimation of trauma rate. (A) Autoregressive time series are random processes where adjacent timepoints are correlated according to *x*_*t*_ = *cx*_*t*−l_ + *𝒩*(0,0.1). The consequence of this is that the random attack rate *x* is correlated across longer timescales, depending on the value of c. Timescales of correlation are shown for five values of c, from lowest (dark blue) to highest (light blue). (B) Example attack rates produced by such an autoregressive time series. (C) These example attack rates can then be used to produce attack sequences for each c value, which can enable analysis of the performance of an optimal Bayesian agent. For n=10000 simulations per autocorrelation (c) value, the error of estimated attack rates is highest for long timescales of autocorrelation.

To test this idea, we induced stress in a mouse model of PTSD (Stress-Enhanced Fear Learning; SEFL) and compared the performance of temporal difference learning (RL model) and a momentum model (RL momentum model) in explaining defensive behavior (Figure 5). In this model, mice receive unpredictable footshocks in one context (Context A) and then show sensitized threat responses to a single footshock in another context (Context B) later (Figure 5A, top). The RL momentum model fits the observed freezing behavior (Figure 5A, bottom) well, showing a disproportionate freezing response to the single footshock in a novel context. This sensitized freezing behavior can be explained by the momentum term in the model, which links the threat prediction errors produced across contexts.

**Figure 5.**
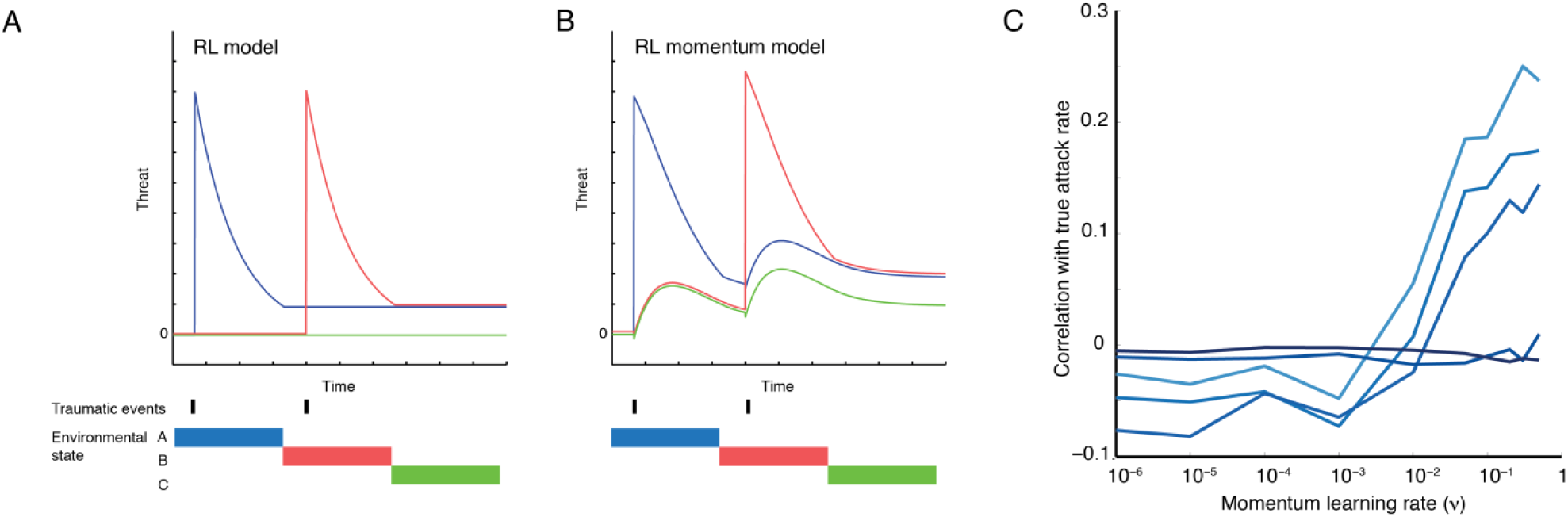
Reinforcement learning with momentum allows improved estimation of autocorrelated attack rates. (A) Single traumatic events occur in different environmental states (contexts), leading to increased associated threat according to the RL model. (B) In the RL momentum model, the same series of attacks produces momentum which couples threat across contexts. Context C threat is due to momentum since the animal receives no footshocks in that state. (C) The momentum learning rate term of the RL momentum model enables extraction of information about fluctuating attack rates. Autoregressive attack rates were produced as shown in figure 3 to produce n=10000 simulated attack sequences (light blue, highest autoregression to dark blue, lowest autoregression). All attacks occur in a different context. In the absence of momentum, the agent cannot extract information about fluctuations in underlying attack rate. With higher momentum, the agent can extract information about the underlying attack rate fluctuations.

We compared Maximum Likelihood fits between the RL and RL momentum models (n=18 unstressed, n=17 stressed mice), using the Bayes Information Criteria (BIC; Figure 5B). When the momentum learning parameter (*ν*) is zero, the two models are equivalent, but the the RL momentum model has a greater number of parameters (4 for RL momentum, 2 for RL model). Since the BIC penalizes the number of parameters, this produces model fits where the RL model is preferred (for unstressed mice, RL model was preferred in 17/18 animals). For stressed mice, however, the BIC strongly favored fits from the RL momentum model (14/17 animals). The RL momentum model predicts greater freezing in a novel context in stressed animals than the RL model, which accounts for the improved predictions over the RL model.

The RL-momentum model of PTSD presents an additional learning mechanism by which PTSD symptoms may be ameliorated. In the classical RL model of PTSD, extinction learning (Figure 6A) works to reduce PTSD by generating small prediction errors when the agent is re-exposed to the traumatic context. This approach underlies evidence-based psychological therapies for PTSD, such as prolonged exposure and cognitive reprocessing therapy. The RL momentum model retains extinction of learned associations, but the threat prediction errors generated by extinction also generate negative momentum that reduces responses to novel threats (Figure 6B). This model also offers a novel perspective on treatment failure of exposure therapy in PTSD.

**Figure 6.**
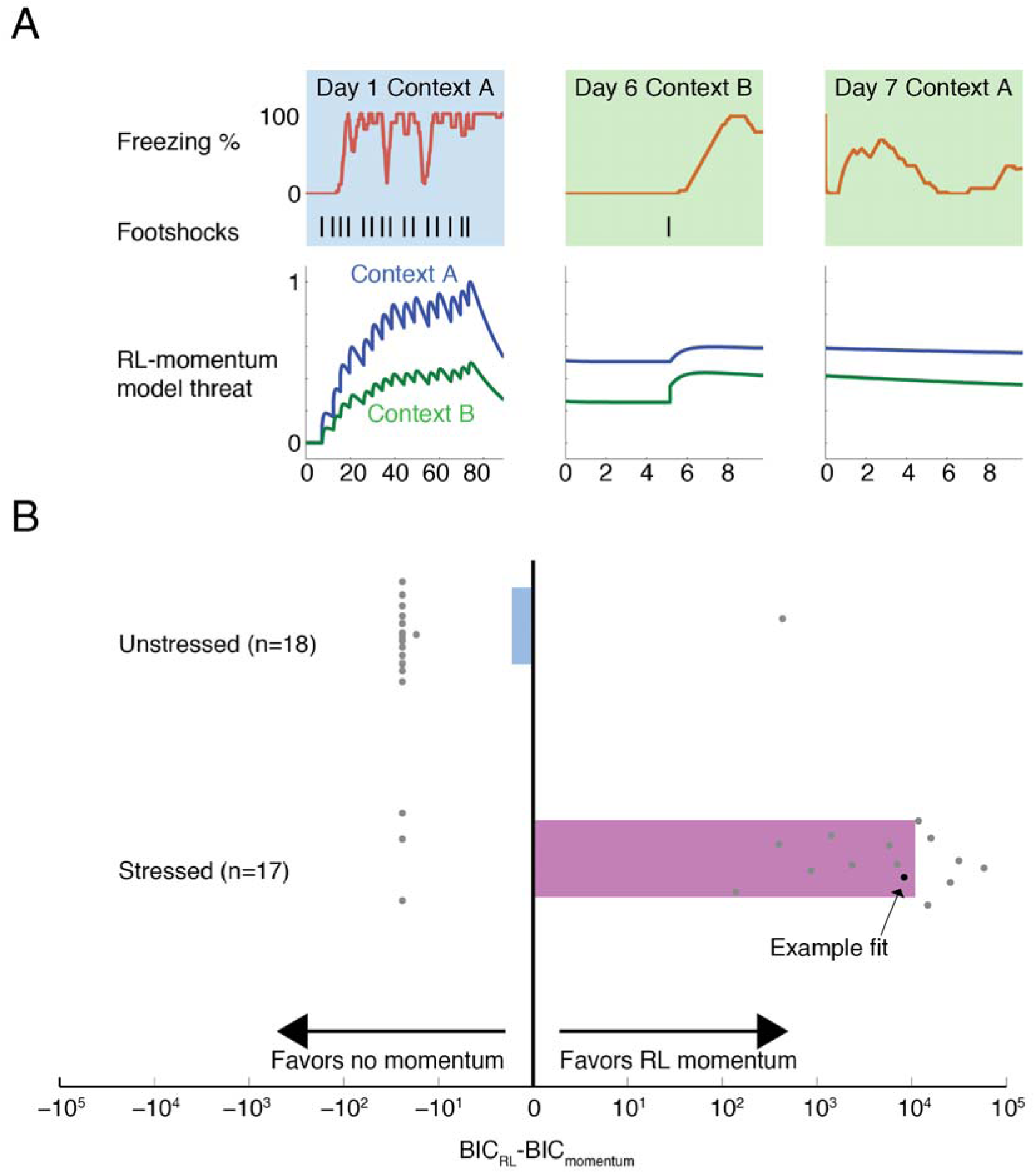
RL momentum fits threat behavioral data in a mouse model of PTSD. (A) Example mouse behavioral data across three days of in the stress-enhanced fear learning model of PTSD (upper), along with RL momentum fit to behavioral data (lower). (upper left) Freezing across 90 minutes (red) of exposure to 15 unpredictable footshocks (black; 1mA, 1s). (upper center) Freezing across subsequent exposure to 1 uncued footshock in a new context. (upper right) Freezing during re-test in the new context (lower left) Threat according to maximum likelihood model fit of the RL momentum model (threat associated with context A – blue, context B-green) on day 1, (lower center) day 6, and (lower right) day7. (B) Model comparison between classic RL model and RL momentum model for SEFL mice (n=17 stressed, n=18 controls). Bayes information criterion (BIC) was calculated (see Methods) for maximum likelihood fits of the RL model and RL momentum model for either unstressed animals (0 shocks on day 1) or stressed animals (15 shocks on day 1). Difference in BIC between the two models is shown for individual animals (gray dots; black dot for example data from (A)), mean BIC difference per condition as bars (blue – unstressed, pink – stressed).

**Figure 7.**
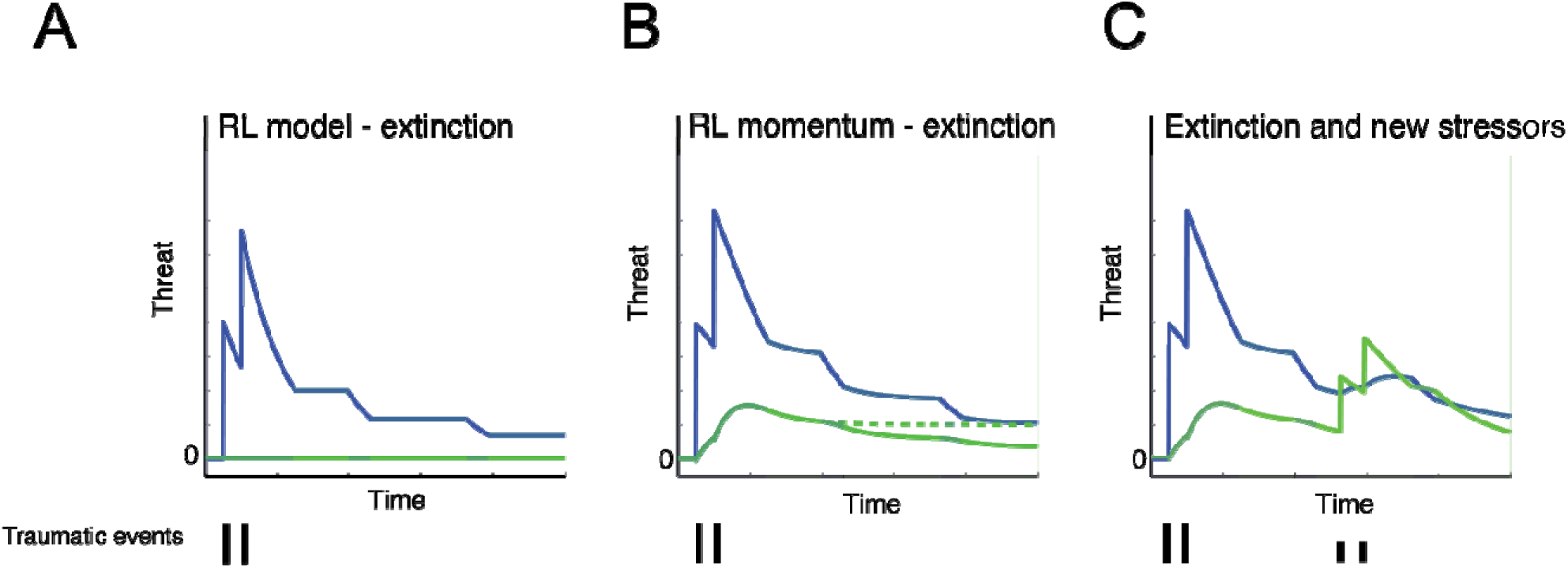
RL momentum model offers a new perspective on mechanisms of extinction and symptom exacerbation in PTSD. (A) RL model: Two traumatic events in an initial context (context A; blue highlight) produce threat learning associated with that context (blue line) but no threat associated with a novel contet (context B; green line) during exposure to that context (green highlights). Extinction occurs when exposure to the initial context A after the traumatic events causes threat prediction errors which decrease threat associated with context A (blue highlights, second and third exposures). (B) RL momentum model: Two traumatic events in initial context produce a momentum which increases threat in a novel context (green line). Re-exposure to initial threat context (context A; blue highlights) reduces threat associated with context A (blue line) but also reduces threat momentum (green line). Green dotted line shows counterfactual threat momentum if no re-exposure to context A had occurred). (C) RL momentum model demonstrates a novel explanation for relapse during exposure therapy. Exposure to smaller stressors (small lines) in a novel context increases threat associated with context B (green line) but also, via the momentum term, increases threat associated with the initial traumatic context A (blue line).

Current learning-based accounts of this phenomenon posit that individuals may experience extinction renewal or extinction resistance, in which either extinction fails to occur or in which the extinction memory may be specific to the context in which it was generated (e.g., the therapy session). In contrast, the RL momentum proffers a simple explanation – unrelated mild stressors generate threat momentum, which increases threat associated with the original traumatic context (Figure 6C). Similarly, an implication of this model is that exposure to novel threats independent of the traumatic context could reduce threat momentum. For example, an agent encountering an intense innate threat (e.g., standing on the side of a high cliff) without injury might experience a strong negative prediction error which would reduce threat momentum for the same reason as exposure to a cue associated to a traumatic event.

## Discussion

We formulated PTSD as a learning process directed at estimating the rate of trauma rather than the specific associations with the trauma. The Bayesian formulation of this problem treated the agent experiencing trauma as an ideal observer. We found that the rate of traumatic events could be estimated well by this agent. Early life trauma had disproportionate impact in this model even without specialized mechanisms for amplifying early life experience. We applied the reinforcement learning momentum model to PTSD, and found that RL-momentum performs well when violence is clustered in time. The slower the change in trauma rate, the more momentum contributes to optimal learning from traumatic stress. This model also offers a novel conceptualization of extinction learning, and suggests that exposure to unassociated strong threats could affect threat momentum. Understanding the impact of innate danger on threat momentum requires further modeling and empirical investigation, since exposure to innate threat could lead to either positive or negative changes in threat momentum.

Previous formal approaches to learning in PTSD have focused primarily on associative mechanisms. However, experimental observations of sensitization to new threats by previous stress are often used to model PTSD (26,29-30). We show that stress sensitization of threat, a model of PTSD, is well fit by the RL-momentum model. However, our ability to precisely fit the parameters of the RL-momentum model is limited by the binary nature of the stress in this dataset. Full validation and parameter-fitting for the RL-momentum model will require more precise manipulations of the sequence of threat prediction errors over time.

A further limitation of this study is that we did not consider parameter regimes that may give rise to habituation (decrease in response to repeated stimuli). Both sensitization and habituation can occur in the RL-momentum model, depending on chosen parameters (7). In PTSD, habituation has recently been suggested as an outcome of repeated trauma (44), and may relate to the numbing symptoms in PTSD. Habituation and sensitization have been thought of as separate processes which competitively modulate responses to repeated stimuli (45). PTSD involves both excessive (hyperarousal) and decreased (numbing) emotional reactions occur after traumatic stress (45-47). A more complete model of the impact of a sequence of threat prediction errors on subsequent emotional responses may explain this apparent contradiction.

Future progress in understanding the role of non-associative learning in PTSD may depend on measuring the neural substrate of threat momentum (or estimated attack rate in the Bayesian model). Applying David Marr’s three levels of analysis to non-associative learning from threat (Figure 1), we have defined the computational problem (“predicting future threats based on a sequence of attacks”) that must be solved. We have compared two algorithms for accomplishing this goal: Bayesian MCMC sampling and RL momentum. We find the RL momentum model offers a formal mathematical approach at the implementation level which explains clinical features of PTSD and behavior in a mouse model of PTSD. However, the implementation level of the RL momentum has not been identified.

Identifying PTSD with threat momentum may facilitate future neurobiological and translational studies of PTSD. Extensive work has shown that patients with PTSD have different learning rate parameters during fear and extinction learning (11-15) than controls in the formation of associations. This study extends these findings by offering a model of how the sequence of threat prediction errors may generate other associative learning alterations in PTSD. The neurobiological correlates of threat momentum would be slowly varying summing functions of previous threat prediction errors which sensitize defensive behaviors, such as neuromodulatory systems (29) or molecular switches leading to persistent neural changes (48). Future extensions of this approach may link effects of arousal on learning rates (rather than overall threat) to averaged recent threat prediction errors, similar to Pearce-Hall learning (49). Thus, the present study may facilitate future work linking non-associative and associative mechanisms in PTSD. Such links are evident in behavioral and epidemiological data and have plausible biological mechanisms, but have previously lacked a computational model to facilitate the design of future experiments.

